# Single-locus species delimitation and a new phylogeny for the genus *Virilastacus*

**DOI:** 10.1101/2025.06.30.662448

**Authors:** Luis Amador, Pedro F. Victoriano, Christian Muñoz-Escobar, Guillermo D’Elía

**Author notes:** Corresponding author: Guillermo D’Elía.

## Abstract

South American crayfish of the family Parastacidae inhabit freshwater systems in northern Argentina, southeastern Brazil, central and southern Chile, and Uruguay. The Chilean endemic genus *Virilastacus*, one of the three South American genera, currently comprises four species distributed across southern Chile. However, the genetic variation and population distribution of *Virilastacus* remain poorly understood, and species delimitation has made limited use of genetic data. In this study, we analyzed mitochondrial DNA sequences to investigate the genetic and geographic species boundaries of *Virilastacus* from a molecular perspective. Divergence time estimates indicate that the stem age of *Virilastacus* is ancient, with the radiation of extant species beginning in the Middle Oligocene. Single-locus species delimitation analyses support five species: the four currently recognized species and a candidate species identified in this study (*V. sp. ‘Calfuco’*). The taxonomic distinction of this candidate species should be further evaluated using additional evidence. Furthermore, our findings suggest an expanded range for *V. jarai* and a reduced distribution for *V. araucanius*. Finally, we discuss the broad geographic patterns of genetic diversity in *Virilastacus* in the context of orographic and paleoclimatic factors.

## Introduction

Defining species boundaries is a fundamental goal in organismal biology and essential for understanding the origins and maintenance of biodiversity. Species are the basic biological unit in ecology, evolutionary biology, biogeography, and conservation biology (Riddle & Hafner, 1999; Cracraft, 2002), making their accurate delimitation imperative. However, this task is particularly challenging for morphologically conservative groups (Hillis, 1987), where cryptic evolutionary lineages and their genetic diversity risk being overlooked. The failure to identify cryptic species can be exacerbated under climate change scenarios, which can obscure biodiversity loss by underestimating the effects of geographic range reductions (Bálint et al., 2011). Despite these challenges, overcoming them is crucial. Effective species delimitation requires integrating diverse evidence, including morphological, genetic, and ecological data (Rissler & Apodaca, 2007). DNA-based methods offer a rapid and effective starting point for identifying candidate species (Burbrink & Ruane, 2021). Examples include studies on frogs (Amador et al., 2018; Jadin et al., 2021) and crayfish (Miranda et al., 2018; Amador et al., 2021).

Currently, many taxonomic groups are experiencing an era of species discovery (e.g., D’Elía et al., 2019; Rouhan & Gaudel, 2021), driven by increased access to remote regions and advanced technologies for species identification and description. The decapod family Parastacidae exemplifies this trend (e.g., Huber et al., 2018; McCormack & Raadik, 2021; Patoka et al., 2023; Ribeiro & Araujo, 2024). For example, within the genus *Parastacus*, 16 new species have been described over the past decade (e.g., Ribeiro et al., 2016, 2017; Huber et al., 2020, 2024a, 2024b). This family of freshwater crayfish, characterized by an ancient and complex evolutionary history, originated in Gondwana (Toon et al., 2010) and is now distributed across Madagascar, Australia, New Zealand, New Guinea, and South America. However, many of the more than 200 species in this family face significant threats, primarily due to habitat loss driven by anthropogenic pressures (e.g., urbanization, stream channelization) and the impacts of climate change (Richman et al., 2015).

In South America, three extant genera (*Parastacus*, *Samastacus*, and *Virilastacus*) are found in distinct regions: northern Argentina (*Parastacus* and *Samastacus*), southern Brazil (*Parastacus*), central and southern Chile (*Parastacus*, *Samastacus*, and *Virilastacus*), and Uruguay (*Parastacus*). Among these, *Virilastacus* is endemic to Chile and comprises four species distributed across southern Chile, inhabiting marshland areas from the Ñuble Region (∼36°S) to the Los Lagos Region (∼42°S). Notably, three of the four species were described within the past two decades (Rudolph & Crandall, 2005, 2007, 2012), reflecting the recent progress in understanding this genus. However, geographic distributions of *Virilastacus* species remain poorly characterized due to limited surveys and large areas within their known ranges remain unsampled. Similarly, little is known about their genetic variation, with the only information about the genetics and morphology of *Virilastacus* coming from species description articles.

To address gaps in knowledge and evaluate the taxonomy of *Virilastacus*, we used mitochondrial DNA (mtDNA) variation to assess the diversity and species limits of crayfish species within the genus throughout its known geographic range. We applied the General Lineage Concept of Species (de Queiroz, 1998), where species are defined as segments of population-level (metapopulation) lineages and multiple operational criteria and lines of evidence may be used to document them. With this work, we hope to contribute to the knowledge of the species diversity within the genus *Virilastacus* and promote new conservation and ecological research.

## Materials and Methods

### Study region and field sampling

*Virilastacus* samples were collected while collecting target specimens of *Parastacus nicoleti* and *P. pugnax* in the Ñuble, Biobío, Araucanía, Los Ríos and Los Lagos regions, sampling was made with special emphasis on covering all major watersheds in the study area (36° 28′ S, 72° 13′ W — 41° 25′ S, 73° 46′ W; 24 sampling localities; Fig. 1). Geographic coordinates, altitude, and habitat data were recorded at each sampling location. A PVC-made manual vacuum pump was used to extract the animals from their subterranean galleries. The specimens were taken to the laboratory and then sacrificed by cryoanesthesia, labeled, and preserved in 96% ethanol. Muscle tissue from one of the walking pereiopods and/or one of the chelipeds was extracted from each of the collected specimens. Voucher specimens were lodged at Departamento de Zoología and Instituto de Ciencias Ambientales y Evolutivas collections in the Universidad de Concepción and Universidad Austral de Chile, respectively.

**Fig. 1.**
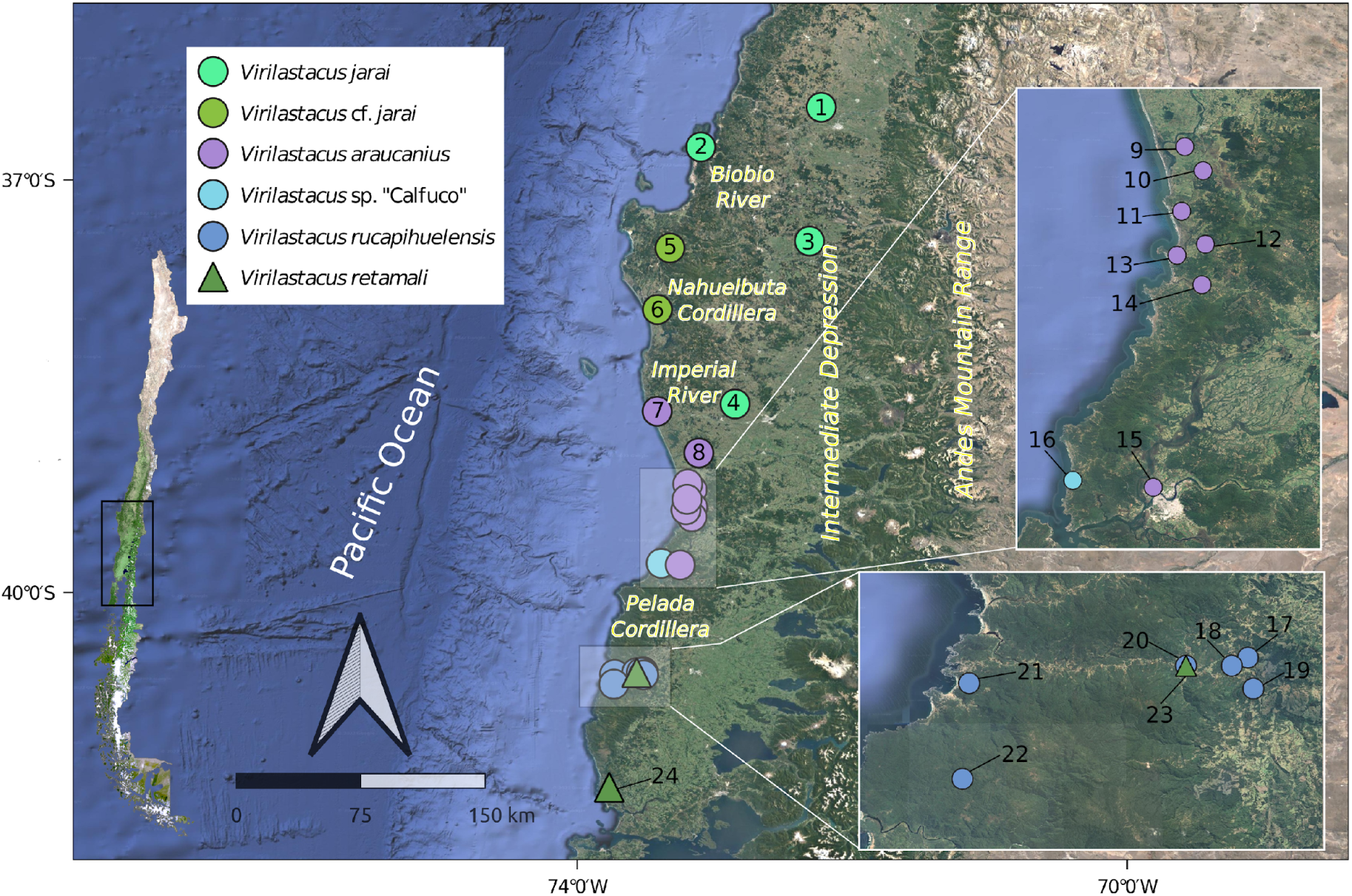
Map of central-southern Chile showing the sampling localities of *Virilastacus* with DNA sequences and the lineages identified in this study (numbers correspond to localities in Table 1).

**Table 1.** Collection localities of the specimens of *Virilastacus* used in this study. Each locality name includes a number that corresponds to those mapped in Fig. 1. For each locality, the catalog number of the analyzed specimens (Genbank accession numbers of the sequences of each specimen are given between parenthesis), geographic coordinates, and the source of the sequence data are indicated.

| Species | Locality | Specimens | Latitude | Longitude | Source |
| --- | --- | --- | --- | --- | --- |
| *Virilastacus jarai* | San Nicolas [1] | 1259SN (OR016100),<br>1261SN (OR016101) | -36.471 | -72.225 | This study |
| *Virilastacus jarai* | Denavi Sur [2] | CC356DS (OR016122) | -36.76 | -73.101 | This study |
| *Virilastacus jarai* | Los Angeles [3] | KC4429a (JQ844466),<br>KC4429b (JQ844467),<br>KC4429c (JQ844468) | -37.444 | -72.31 | Rudolph and<br>Crandall, 2012 |
| *Virilastacus jarai* | Cholchol [4] | CC672CH (OR016102),<br>CC676CH (OR016104),<br>CC678CH (OR016105),<br>CC679CH (OR016107),<br>CC680CH (OR016106),<br>CC681CH (OR016103) | -38.629 | -72.851 | This study |
| *Virilastacus* cf.<br>*jarai* | Curanilahue Sur<br>[5] | CC758CUR (OR016109),<br>CC759CUR (OR016111),<br>CC760CUR (OR016108),<br>CC761CUR (OR016110) | -37.495 | -73.325 | This study |
| *Virilastacus* cf.<br>*jarai* | Colo Colo [6] | CC838CC (OR016120),<br>CC839CC (OR016113),<br>CC840CC (OR016117),<br>CC841CC (OR016119),<br>CC842CC (OR016118),<br>CC843CC (OR016115),<br>CC844CC (OR016116),<br>CC845CC (OR016114),<br>CC846CC (OR016121),<br>CC847CC (OR016112) | -37.942 | -73.416 | This study |
| *Virilastacus*<br>*araucanius* | Fundo Chomio<br>[7] | 2219FC (OR016065),<br>2220FC (OR016066) | -38.682 | -73.418 | This study |
| *Virilastacus*<br>*araucanius* | Teodoro<br>Schmidt [8] | PP184 (OR016091),<br>PP185 (OR016090),<br>PP186 (OR016092) | -38.985 | -73.115 | This study |
| *Virilastacus*<br>*araucanius* | La Barra [9] | CC864LB (OR016093),<br>CC865LB (OR016075),<br>CC866LB (OR016067),<br>CC867LB (OR016079),<br>CC868LB (OR016081),<br>CC869LB (OR016080),<br>CC870LB (OR016076),<br>CC871LB (OR016078), | -39.21133<br>3 | -73.195583 | This study |
|  |  | CC872LB (OR016077),<br>CC873LB (OR016074) |  |  |  |
| *Virilastacus*<br>*araucanius* | Chanquin [10] | PP179 (OR016088),<br>PP180 (OR016089),<br>PP181 (OR016085),<br>PP182 (OR016086),<br>PP183 (OR016087) | -39.253 | -73.164 | This study |
| *Virilastacus*<br>*araucanius* | Between Tolten<br>and Queule [11] | PP187 (OR016070),<br>PP188 (OR016072),<br>PP189 (OR016094),<br>PP190 (OR016069) | -39.323 | -73.201 | This study |
| *Virilastacus*<br>*araucanius* | Piren Bajo [12] | PP267 (OR016096),<br>PP268 (OR016099),<br>PP269 (OR016098),<br>PP270 (OR016097) | -39.38 | -73.16 | This study |
| *Virilastacus*<br>*araucanius* | Queule [13] | PP174 (OR016084),<br>PP175 (OR016095),<br>PP176 (OR016071),<br>PP177 (OR016068),<br>PP178 (OR016073) | -39.399 | -73.209 | This study |
| *Virilastacus*<br>*araucanius* | Piutril [14] | PP168 (OR016083),<br>PP170 (OR016082) | -39.45 | -73.166 | This study |
| *Virilastacus*<br>*araucanius* | Universidad<br>Austral [15] | KC1416 (EF599156) | -39.8 | -73.25 | Crandall et al.,<br>2000 |
| *Virilastacus* sp.<br>'Calfuco' | Calfuco [16] | PP322 (OR016064) | -39.788 | -73.389 | This study |
| *Virilastacus*<br>*rucapihuelensis* | Carrico [17] | PP403 (OR016062),<br>PP404 (OR016060) | -40.577 | -73.521 | This study |
| *Virilastacus*<br>*rucapihuelensis* | Coigueria [18] | -- | -40.583 | -73.533 | Rudolph and<br>Crandall, 2005 |
| *Virilastacus*<br>*rucapihuelensis* | Contaco [19] | -- | -40.6 | -73.517 | Rudolph and<br>Crandall, 2005 |
| *Virilastacus*<br>*rucapihuelensis* | Rucapihuel [20] | KC2679 (EF599148),<br>KC2680 (EF599149),<br>KC2681 (EF599150) | -40.583 | -73.567 | Rudolph and<br>Crandall, 2005 |
| *Virilastacus*<br>*rucapihuelensis* | Maicolpue [21] | PP333 (OR016061),<br>PP334 (OR016063) | -40.596 | -73.728 | This study |
| *Virilastacus*<br>*rucapihuelensis* | Loma de la<br>Piedra [22] | -- | -40.667 | -73.733 | Rudolph and<br>Crandall, 2005 |
| *Virilastacus*<br>*retamali* | Rucapihuel [23] | KC2684 (EF599153),<br>KC2685 (EF599152),<br>KC2686 (EF599151),<br>KC2687 (EF599154),<br>KC2688 (EF599155) | -40.583 | -73.567 | Rudolph and<br>Crandall, 2007 |
| *Virilastacus*<br>*retamali* | Estaquilla [24] | -- | -41.417 | -73.767 | Rudolph and<br>Crandall, 2007 |

### DNA data acquisition and sequencing

We analyzed 75 sequences of 630 base pairs of the mitochondrial gene (mtDNA) cytochrome c oxidase subunit 1 (COI) from specimens representing the four species of *Virilastacus* gathered at 11 localities and covering the distribution range of the genus (Fig. 1, Table 1). Twelve of these sequences were downloaded from GenBank, while the other 63 sequences were gathered here following the protocol outlined below. To set the outgroup, we included 15 sequences of different species of Parastacidae available in GenBank including seven of *Parastacus* and one of *Samastacus spinifrons* as well as seven species from Madagascar, Australia, and New Zealand. GenBank accession numbers of new sequences (OR016060 – OR016122) and for all other sequences included in the analyses are provided in Table S1.

DNA extractions were done using the commercial kit Wizard SV genomic (Promega) following the manufacture protocol. A fragment of the COI mtDNA gene was amplified via PCR using the primers provided by Folmer et al. (1994) with the following conditions: initial denaturation at 94 °C for 3 minutes followed by 35 cycles of denaturation at 94 °C for 45 seconds, annealing at 48 °C for 30 seconds; and extension (elongation) at 72 °C for 1:30 minutes and a final extension step at 72 °C for 10 minutes. Amplicons were purified and sequenced at Sequencing Group Macrogen Inc., Korea. Electropherograms were visualized and edited using Geneious (Biomatters Ltd., USA); a visual inspection searching for internal codon stops and reading frame shifts was performed. All new sequences were deposited in GenBank (accession numbers in Table S1).

### Sequence alignment and matrix reconstruction

Sequences were aligned with MAFFT v7 (Katoh & Standley, 2013) as implemented in Geneious Prime 2 (https://www.geneious.com) using default parameter values. The alignment was visualized in AliView (Larsson, 2014) to check for the presence of reading frame shifts. We built three matrices for phylogenetic and species delimitation analyses: 1) a matrix with 90 sequences (ingroup plus outgroup; Supplementary Material) that was used in Maximum Likelihood (ML), Bayesian Inference (BI), and divergence time analyses, 2) a matrix with 14 sequences; six sequences of *Virilastacus* each corresponding to clades identified in the phylogenetic analyses (see below), and eight sequences from the outgroup (seven *Parastacus* and *Samastacus spinifrons*), which was used in analyses of species tree estimation and species delimitation analyses, and 3) a matrix without the sequences of the outgroup, containing the 75 samples of *Virilastacus*, used to calculate genetic distances.

### Phylogenetic analyses and divergence time estimate

The best-fit DNA substitution model for the first two matrices was obtained using ModelFinder (Kalyaanamoorthy et al., 2017) and selected according to the Bayesian Information Criterion (BIC), we did the model selection allowing comparison of different substitution models using codon positions in alignments. Phylogenetic analyses were performed using both ML and BI. ML analyses were conducted using IQ-TREE 2 (Minh et al., 2020), with default values for tree search parameters (i.e., number of initial parsimony trees = 100, number of top initial parsimony trees to optimize with ML nearest neighbor interchange (NNI) to initialize the putative set = 20, number of trees in the putative set to maintain during ML tree search = 5, number of unsuccessful iterations to stop = 100). Nodal support was assessed via 1000 replicates of ultrafast bootstrapping (UFBoot2; Hoang et al., 2018). We used BEAST2 v. 2.4.7 (Bouckaert et al., 2014) to infer Bayesian gene trees for each matrix, using the corresponding best-fit substitution model, a relaxed clock model, and a birth-death tree prior. A run of 5 x 10^7^ generations sampled every 1000 generations was implemented in each analysis. A time calibrated-ultrametric tree was also obtained with BEAST2, using as a secondary calibration point the time estimate (i.e., normal distribution with a divergence time estimate = 116 Ma and a confidence interval (95%) = 89 – 142 Ma) for the most recent common ancestor of the South American parastacids as inferred by Toon et al. (2010). Analysis used a relaxed clock log normal model and a Calibrated Yule model of speciation. Runs lasted for 50 x 10^7^ generations and were sampled every 1,000 generations. Trace log files were checked for convergence and ESS values above 200 using Tracer 1.7 (Rambaut et al., 2018), maximum clade credibility trees, with posterior probability (PP) values, were estimated with the program TreeAnnotator (distributed as part of BEAST2) with the sampled trees after discarding the first 25% as burn-in. All phylogenetic trees were visualized using FigTree 1.4 (Rambaut, 2014).

### Estimates of genetic distance

We used MEGA 11 (Tamura et al., 2021) to estimate divergence values (p-distances) between pairs of individual sequences and within and between pairs of clades identified in phylogenetic analyses (see below); positions with less than 95% site coverage were eliminated (partial deletion option). The robustness of the estimates was assessed with 1000 bootstrap replicates.

### Species delimitation analysis

Single-locus species delimitation analyses were conducted with four different methods based on the phylogenetic trees and genetic distances as inferred above. We first used the Generalized Mixed Yule Coalescent (GMYC; Pons et al., 2006; Fujisawa & Barraclough, 2013) that tries to identify the transition between a coalescent process (population-level) and a Yule process (species-level) was implemented in the R package *splits* (Ezard et al., 2009) using simple and multiple threshold models (GMYCs, Pons et al., 2006; GMYCm, Monaghan et al., 2009), using the ultrametric tree obtained with BEAST2 as input. The same guide tree was used to implement, with the R package *bGMYC* (Reid & Carstens, 2012), the Bayesian version of GMYC (bGMYC), which calculates the marginal posterior probabilities of species limits from the posterior distribution of ultrametric trees. For the bGMYC analysis, the priors of parameters t1 and t2 were set at 2 and 15, respectively. The bGMYC analysis was performed with 100,000 generations, with a burn-in of 50%, and a thinning interval of 10 samples. We also used PTP (Poisson tree process; Zhang et al., 2013) that model speciation and coalescence events in terms of the number of substitutions providing a hypothesis of species limits based on a phylogenetic tree trying to identify the most likely classification of branches at the population and species levels. Additionally, we implemented two recently developed methods of species delimitation. The first one was Assemble Species by Automatic Partitioning (ASAP; Puillandre et al., 2021), which proposes partitions of species hypotheses using the genetic distances calculated among DNA sequences. Finally, we used DELINEATE (Sukumaran et al., 2021) which introduces the framework ‘speciation-based delimitation’ to distinguish species and population boundaries using probabilistic models, through the incorporation of an extended speciation process model into the delimitation analysis. For DELINEATE we used as input the ultrametric tree inferred with BEAST2.

### Morphological analysis

To confirm the identity of species recovered on the phylogenetic analysis, we performed a single qualitative morphological analysis in specimens collected in the locality of Maicolpue (locality 21), which were grouped in the same clade with *Virilastacus rucapihuelensis* and where *V. araucanius* was reported by Bahamonde and collaborators in 1998 (Rudolph, 2015). We followed the original description of *V. rucapihuelensis* (Rudolph & Crandall, 2005) and the characters used to distinguish it from other *Virilastacus* species.

## Results

### Phylogenetic analyses and molecular divergence

The resulting phylogeny recovered *Virilastacus* monophyletic (uBS = 77; PP = 93) and sister to *Samastacus* with robust support values (uBS = 83; PP = 0.98). Within *Virilastacus*, each of the four recognized species was also recovered monophyletic. The basal dichotomy of the clade of *Virilastacus* leads on one hand to *V. jarai* (uBS = 100; PP = 0.98). This clade shows geographic structure; there are two latitudinal allopatric subclades (Figs. 2, S1). One is composed of variants from two localities to the north of Biobío River and two localities to the east of Nahuelbuta Cordillera. The other subclade is represented by variants from two localities distributed in the western part of the Nahuelbuta Cordillera (Fig. 2). The second main clade of *Virilastacus* showed no support and is composed of sequences gathered from all other individuals of the genus, including *V*. *araucanius*, *V*. *rucapihuelensis*, *V*. *retamali*, and from a single divergent individual of the coastal locality of Calfuco in Los Rios Region (Fig. 1). *V*. *rucapihuelensis* and *V*. *retamali*, distributed in the southernmost range of genus *Virilastacus* in the Los Lagos Region (Fig. 1), are recovered as sister species both in ML and BI analyses (uBS = 85; PP = 0.97; Figs 2, S1). In the ML tree, the sample from Calfuco forms a weakly supported clade with *V*. *rucapihuelensis* and *V*. *retamali* (uBS = 76; Fig. S1). This latter clade collapse in the BI analysis appears as a trichotomy involving *V. araucanius*, the single sequence from Calfuco, and the clade composed by *V*. *rucapihuelensis* and *V*. *retamali*.

**Fig. 2.**
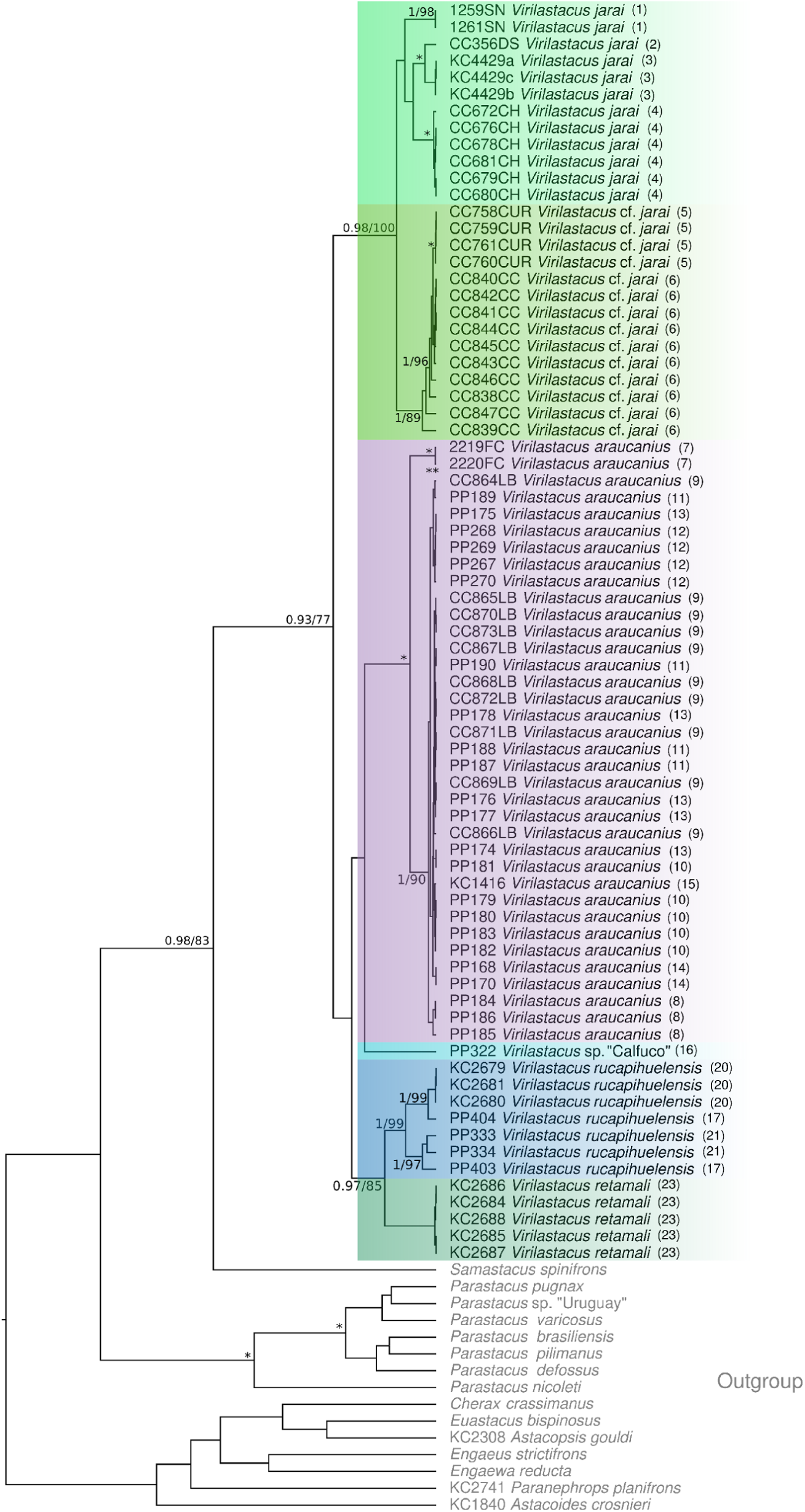
Bayesian tree showing the relationships of species of *Virilastacus* for the COI data. Numbers in nodes correspond to posterior probability (Bayesian inference) / bootstrap support (Maximum Likelihood) values. Asterisks indicate perfect support for that node (* = 1/100). The colors correspond to species localities in Fig. 1.

### Genetic distances

The average of uncorrected p-distances between the six main lineages (i.e., the four recognized species of *Virilastacus* species and the two other identified lineages, *V*. cf. *jarai* and *V*. sp. ‘Calfuco’) range from 0.077 (*V*. *jarai s.s.* vs. *V*. cf. *jarai*; Table S2) to 0.142 (*V*. *araucanius* vs. *V*. *rucapihuelensis*; Table 2). Comparisons involving *V*. *jarai* – cf. *jarai* as a single species have p-distances from 0.122 to 0.124; while those involving *V*. sp. ‘Calfuco’ goes from 0.105 to 0.122 (Table 2). Average comparisons within clades present the divergence within *V*. *rucapihuelensis* as the highest value (0.050) and *V*. *retamali* presents the lowest value (0.000) (Table S3).

**Table 2.** Average genetic distances between pairs of five lineages of *Virilastacus*, following the five-species scenario proposed in species delimitation analyses.

|  | *V. rucapihuelensis* | *V. sp. 'Calfuco'* | *V. retamali* | *V. araucanius* |
| --- | --- | --- | --- | --- |
| *V. rucapihuelensis* |  |  |  |  |
| *V. sp. 'Calfuco'* | 0.105 |  |  |  |
| *V. retamali* | 0.100 | 0.111 |  |  |
| *V. araucanius* | 0.142 | 0.118 | 0.116 |  |
| *V. jarai* (including cf. *jarai*) | 0.124 | 0.122 | 0.122 | 0.122 |

### Species delimitation

Single and multiple thresholds in GMYC analyses (GMYCs and GMYCm respectively) provided different results. GMYCs recovered the six main lineages of *Virilastacus* (i.e., *V*. cf. *jarai*, *V*. *jarai* s.s., *V*. *rucapihuelensis*, *V*. *retamali*, *V*. sp. ‘Calfuco’ and *V*. *araucanius*) as part of a single species, while GMYCm recovered each of the six main lineages as a different candidate species (Fig. 3). The Bayesian implementation of GMYC (bGMYC), recovered four species, *V*. *jarai* s.s. + *V*. cf. *jarai*, *V*. *rucapihuelensis* + *V*. *retamali*, *V*. *araucanius*, and *V*. sp. ‘Calfuco’. This result was also obtained in the first best partition in ASAP analysis (ASAP R1, Fig. 3). Instead, the second-best partition of ASAP (ASAP R2) recovered five species, one formed by *V*. *jarai* s.s. + *V*. cf. *jarai* and the others by each of the remaining four main lineages of *Virilastacus*. PTP gave a scenario of three species, *V*. *jarai* s.s. + *V*. cf. *jarai*, *V*. *rucapihelensis* + *V*. *retamali* +*V*. sp. ‘Calfuco’ and *V*. *araucanius*. Finally, DELINEATE recovered each of the six main lineages as a different candidate species, with the same result as GMYCm.

**Fig. 3.**
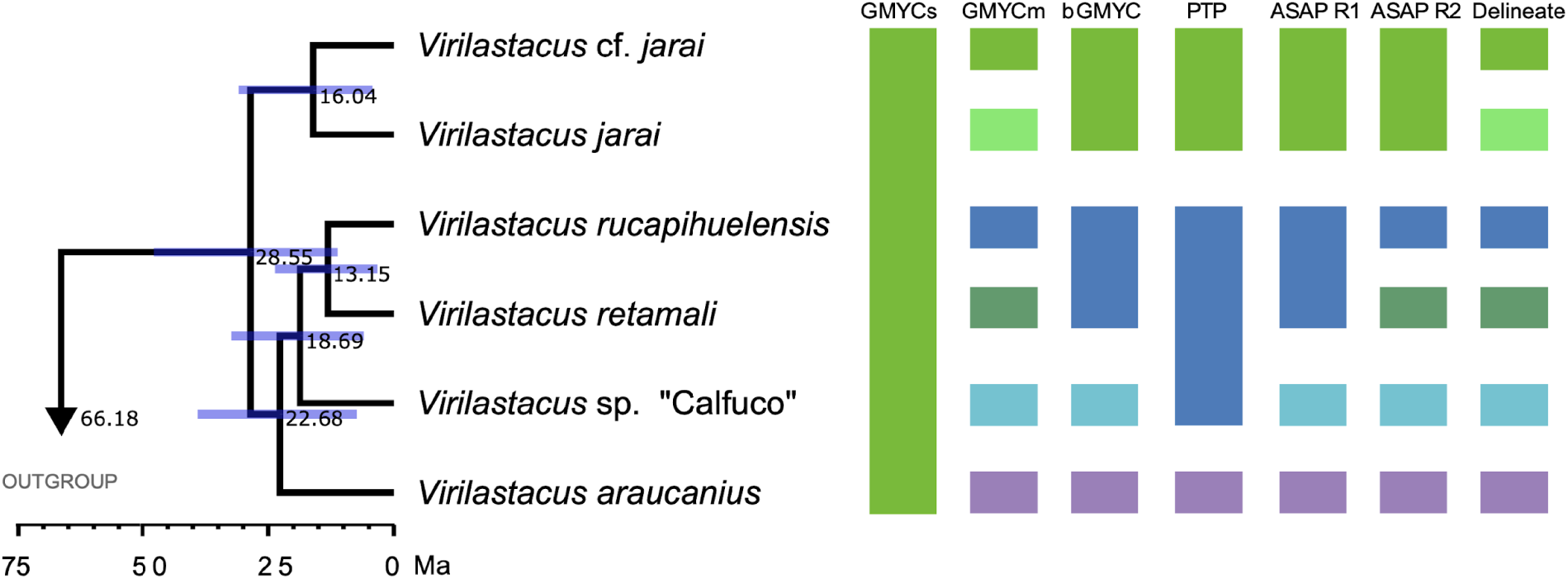
Species tree for *Virilastacus*, with the branches drawn using the posterior means of divergence times. Values in nodes correspond to estimated mean ages in millions of years. Violet bars in the tree indicate 95% HPD intervals. The summary of species delimitation results from seven approaches is shown on the right.

### Divergence times

The split between *Virilastacus* and *Samastacus* was estimated at 66.18 Ma (24.41 – 109.23 Ma). The crown age of *Virilastacus* was dated 28.55 Ma (11.24 – 47.73 Ma) in the Mid-Oligocene. The divergence between the two identified lineages in *V*. *jarai* (*V*. *jarai* s.s. and *V*. cf. *jarai*) was estimated at 16.04 Ma (4.31 – 30.9 Ma). The crown age of the clade formed by the other four main lineages occurred at about 22.68 Ma (7.38 – 39.03 Ma), the split of *V.* sp. ‘Calfuco’ with *V*. *retamali* and *V*. *rucapihuelensis* was 18.69 Ma (6.0 – 32.3) and the split between the latter two occurred between 13.15 Ma (3.33 – 23.65 Ma) (Fig. 3).

### Morphological analysis

Two specimens (PP333 and PP334) were collected in Maicolpue. Based on the following distinctive morphological characteristics: 1) rostral carina long and prominent (Fig. 4A), 2) flap present in the second somite (absent in other *Virilastacus*) (Fig. 4B), 3) telson subrectangular (Fig. 4C), 4) supernumerary gonopores (female or male in other *Virilastacus*) (Fig. 4D), 5) phallic papillae widely separated, robust, and short (Fig. 4D), 6) male cuticle partition present in P5 coxae (absent in other *Virilastacus*) (Fig. 4D). These characteristics were useful to place these specimens as *V. rucapihuelensis*, being concordant with phylogenetic results (Fig. 2).

**Fig. 4.**
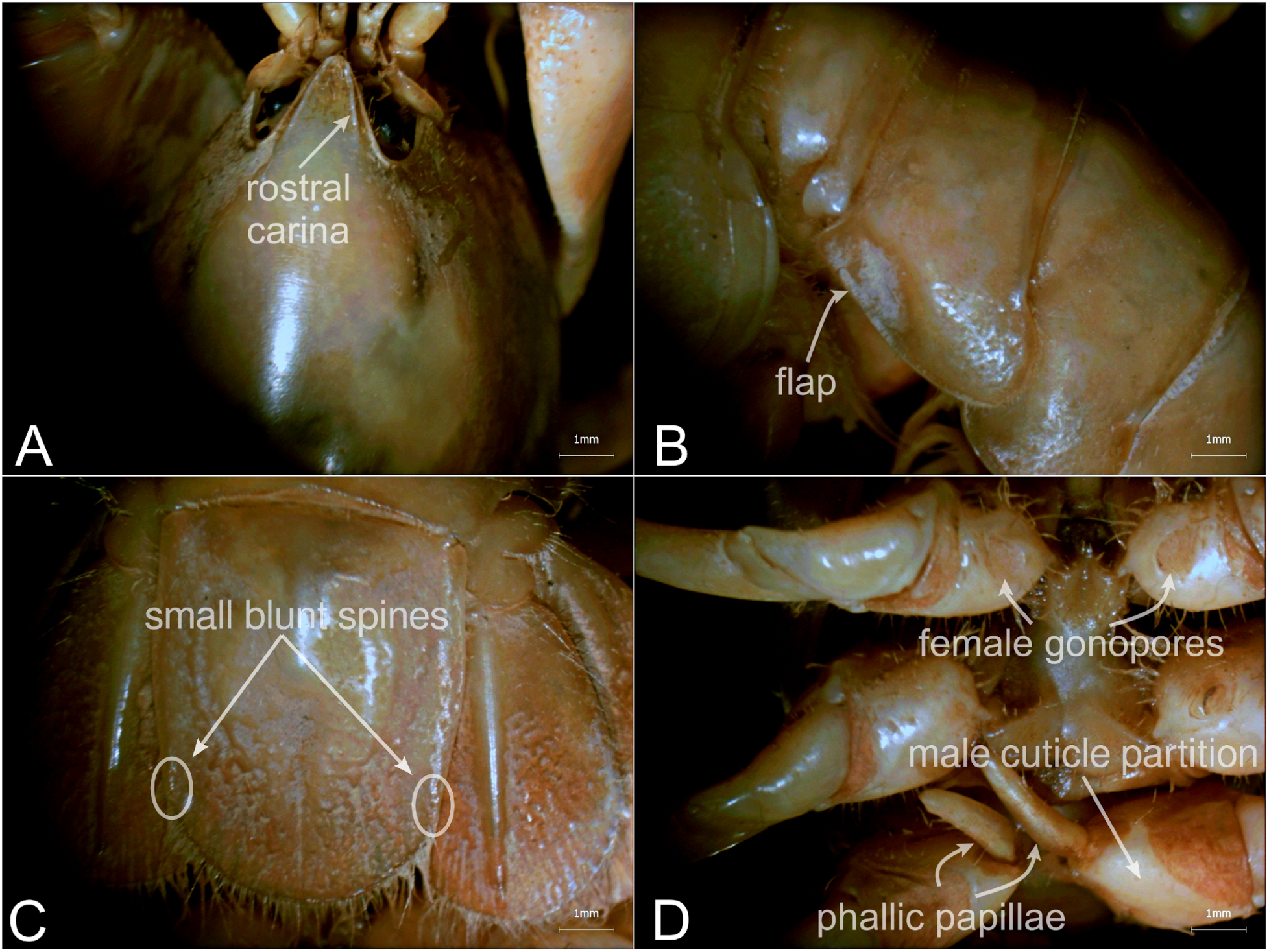
Morphological characters allow the assignment of specimens collected at Maicolpue (locality 21) belonging to *Virilastacus rucapihuelensis*. A: cephalotorax/rostrum, B: pleon/second pleura, C: telson, D: thoracic sternites/gonopores/phallic papillae.

## Discussion

The present contribution is the first study focused on delimiting species based on explicit species discovery methods of the Chilean endemic crayfishes of the genus *Virilastacus* across its geographic distribution. In general, crayfish of *Virilastacus* have been little studied regarding their intraspecific genetic variation. Previous studies have used restricted geographic coverage limited to a few specimens (Crandall et al., 2000; Rudolph and Crandall, 2012).

We found the gene tree geographically structured where its basal split leads to two latitudinal allopatric clades, a northern clade formed by genetic variants of *Virilastacus jarai* s.s. and *V*. cf. *jarai*, and a southern clade formed by the sequences of *V*. *araucanius*, *V*. sp. ‘Calfuco’, *V*. *retamali*, and *V*. *rucapihuelensis* (Figs. 3, S1). As in Rudolph and Crandall (2012), *V*. *jarai* was recovered monophyletic, while its distribution was expanded with more specimens collected at previously non-studied localities. *V*. *jarai* s.s. appears as the sister to a newly uncovered allopatric lineage, *V*. cf. *jarai*, forming a clade that is sister and allopatric to the clade formed by the rest of the species of *Virilastacus* (Figs. 2, 3, S1). The Bayesian Inference analysis in BEAST recovers *V*. sp. ‘Calfuco’ as the sister taxon of *V*. *araucanius*, however, we follow the scenario where *V*. sp. ‘Calfuco’ is the sister taxon of the clade composed by *V*. *retamali* and *V*. *rucapihuelensis*, as recovered by the Maximum Likelihood and species tree analyses. This phylogenetic incongruence reinforces the notion that *V.* sp. ‘Calfuco’ is a different lineage from the other *Virilastacus* species. Future studies would clarify the phylogenetic relationships of the species of *Virilastacus*.

In this work, we used seven approaches for species delimitation, whose results mostly vary. Incongruence among the results of distinct species delimitation methods is a common finding, as is the case of the single previous study implementing these methods with South American parastacids (Amador et al. 2021; see examples in other groups in Amador et al. 2018; García–Melo et al. 2019; Hurtado and D’Elía, 2019). As such, the usage of a single species delimitation method is not recommended (Miralles and Vences, 2013; Luo et al., 2018), being advisable to simultaneously use tree-based models such as PTP, GMYC (bGMYC), and the recent approach implemented in DELINEATE, with models based on genetic distances such as the new program ASAP, as implemented in the present work. We found congruence in species delimitation results in only two methods; GMYCm and DELINEATE were congruent in recovering a scenario of six candidate species, the four currently recognized species of *Virilastacus* plus *V*. cf. *jarai* and *V*. sp. ‘Calfuco’; while bGMYC and ASAP R1 recovered four species, *V*. *araucanius*, *V*. sp. ‘Calfuco’, lumping *V*. *jarai* s.s. with *V*. cf. *jarai*, and *V*. *retamali* with *V*. *rucapihuelensis* (Fig. 3). The latter result could be due to methods based on genetic distances, such as ASAP, which tend to underestimate the number of species. The comparisons between *V*. *jarai* s.s. and *V*. cf. *jarai* (p-distance = 0.077) and *V*. *retamali* and *V*. *rucapihuelensis* (p–distance = 0.10) obtained the lowest values of genetic divergence between pairs of species of *Virilastacus*, which could have prompted each of these species pairs were recovered as the same species with bGMYC and ASAP R1 (*V*. *retamali* + *V*. *rucapihuelensis*), and with bGMYC, PTP, ASAP R1 and ASAP R2 (*V*. *jarai* s.s. + *V*. cf. *jarai*).

Given the considerations above, we propose a conservative scenario of five species, that is, the five-species scenario obtained with the ASAP R2 approach. These five species correspond to the currently four recognized species, *V*. *jarai* (including *V*. cf. *jarai*), *V*. *araucanius*, *V*. *retamali*, and *V*. *rucapihuelensis*, plus a new candidate species, *V*. sp. ‘Calfuco’. Although this five-species scenario was recovered only by one approach, there were areas of agreement among approaches; for example, four of the seven approaches agreed that *V*. *jarai* s.s. and *V*. cf. *jarai* represent a single species; on the other hand, five approaches delimited to *V*. sp. ‘Calfuco’ as a different species. The latter diverges from *V*. *araucanius*, the geographically closest species, by 12.2% (Table 2); the percentage of divergence value is slightly larger than those observed between *V*. *araucanius* and *V*. *retamali* (11.6%), and between *V*. *araucanius* and *V*. *jarai* (11.9%) and is in line with values observed among species of South American parastacids (Amador et al. 2021; Miranda et al. 2018; Victoriano and D’Elía 2021).

It should be noted that *Virilastacus* sp. ‘Calfuco’ was delimited based on a single individual whose preservation condition was poor, hampering the assessment of its morphological distinction. As such, efforts to collect more individuals in the same locality and surroundings are crucial, as well as their analysis based on nuclear, phenotypic, and ecological data to test the distinction of this candidate species. Having said that, we note that even when known from a single locality, this lineage likely would distribute in several small Pacific Ocean coastal areas of the Los Ríos Region mirroring the distribution of a sympatric and still undescribed crayfish species of the genus *Parastacus* (Amador et al., 2021, 2022). However, the fact is that so far, *V*. sp. ‘Calfuco’ is only known from a very small coastal basin of a tiny creek (‘estero’) that flows from nearby hills directly to the Pacific Ocean beach named Calfuco of Los Ríos Region. In turn, this basin lies in an area that is being intensely impacted by urban expansion along the ocean coast. As such, additional surveys are urgently needed to assess the status of this lineage of species level that is already threatened.

Before this work, *Virilastacus jarai* was only known from its type locality (‘El Porvenir’ sector in central-southern Chile, Rudolph and Crandall, 2012; Rudolph, 2015); however, after this study *V. jarai* became the species with the largest known geographic range of all species of *Virilastacus* (∼ 22,100 Km²). We reported five new localities for the species, including two localities (locality 5 – Curanilahue Sur and locality 6 – Colo Colo; Fig. 1, Table 1) that fall within a distributional polygon of *V. araucanius* as portrayed in Rudolph (2015) and De los Rios–Escalante et al. (2016). One of the five new localities of *V*. *jarai* is San Nicolas (locality 1, Fig. 1), which is the easternmost and northernmost known locality of the genus, as such, here we also extend the known distribution of the genus. In addition, with three records, *V*. *jarai* is the only species of the genus that has localities present in the Chilean Intermediate Depression between the Coastal and Andean Cordilleras (Fig. 1), being all other species distributed in the Coastal Cordillera. On the contrary, given our results, the current estimated range of *V*. *araucanius* (∼ 2,805 Km²) decreased by approximately two orders of magnitude. This is due to haplotypes recovered from specimens collected at Maicolpue (locality 21, Fig. 1) are in the same clade as the samples of *V*. *rucapihuelensis* (Fig. 2), confirming that samples of *Virilastacus* from Maicolpue belong to *V*. *rucapihuelensis* and not to *V. araucanius* as previously proposed to decades ago by Rudolph and Crandall (2005). To corroborate this genetic-based result with the morphological analysis of specimens collected in Maicolpue, where *V. araucanius* was reported for the first time (Bahamonde et al., 1998) in a date before the description of the other three *Virilastacus* species. We found that these specimens present the diagnostic characteristics of *V*. *rucapihuelensis* when we compare them with other *Virilastacus* species. For instance, *V. rucapihuelensis* (intersex individual) presents male cuticle partition, absent in the other species of the genus; telson subrectangular with small blunt spines; the presence of a flap in the pleura of the second somite, absent in the other *Virilastacus*; and a shorter, robust and separated phallic papillae (Fig. 4). As such, based on this evidence, we suggest that *V*. *araucanius* should be restricted to a Pacific coastal strip from the Imperial River in the north to the city of Valdivia in the south (Fig. 1).

Divergence times inferred together with the species tree suggest that the period of major diversification in *Virilastacus* was 23–16 Ma, which agrees in part with the acceleration of the uplift of the southern Andes (Boschman, 2021). It is estimated that the relief that established the current watersheds of southern central Chile and that would have delineated the depositional areas that constitute the habitat of *Virilastacus* would have occurred approximately 6 Ma (Farías et al., 2008). The glacial cycles could also prompt the differentiation of the populations of *Virilastacus*; the first glaciations in southern South America started in the late Miocene, about 6 Ma (Rabassa et al., 2011). The latter may have modulated the direction and subsequent fragmentation and isolation of the paleo-basins (Ding et al. 2023). A Miocene diversification has been also suggested for other animal groups from southern Chile, including the species complex of the burrowing crayfish *Parastacus nicoleti* (Amador et al., 2022), ground beetles of the genus *Ceroglosssus* (López-López et al., 2021), and marsupials of the genus *Dromiciops* (Quintero-Galvis et al., 2021). Furthermore, in the northern range of *Virilastacus*, the turnover between *V*. *araucanius* and *V*. *jarai* may be a mixture of relief changes of the Nahuelbuta mountain range, glaciations, and marine transgressions (e.g., Rodríguez Tribaldos et al., 2017) occurred since the Miocene. Nahuelbuta Cordillera has also been invoked as playing a key role in the differentiation in populations of the crayfish *P. pugnax* (Victoriano and D’Elía, 2021) and different taxa including other invertebrates (e.g., red cricket; Alfaro et al., 2018), vertebrates (e.g., lizards; Victoriano et al., 2008), and plants (e.g., monkey puzzle tree; Fuentes et al., 2021). Further analyses need to use distinct molecular markers from those used in the present study; nuclear markers based on massive sequencing (e.g. Single Nucleotide Polymorphisms – SNPs) have been useful in recovering robust relationships within parastacid species (Unmack et al., 2019; Amador et al., 2022). In this context, a complementary source of evidence could be the application of ecological niche models (ENM), from which the present and past potential distribution ranges could be estimated (e.g., Victoriano and D’Elía, 2021). However, the reduced geographical distribution of species of *Virilastacus* and the low number of records known per each species, pose a methodological difficulty in the usage of this approach.

The taxonomy of *Virilastacus* is still in its infancy, and much needed remains to be done before a solid and stable taxonomic framework is attained. Among the tasks that should be pursued, the first one is an extensive collection of specimens both in already known localities as well as in still unsampled areas. These specimen series would allow assessing the congruence between morphology and genetic variation patterns as a way to test the taxonomic scheme advanced here and to better delineate species distributions.

## Conclusion

Our results suggest that the species diversity of *Virilastacus* is underestimated by the current taxonomic scheme of four species. This suggestion is in line with recent scenarios advanced for the Chilean *Parastacus nicoleti* and *P. pugnax* that were robustly shown to constitute species complexes (see Amador et al., 2021, 2022 and Victoriano & D’Elía, 2021) as well as for other animal taxa from the Valdivian ecoregion (e.g., mammals: D’Elía et al. 2015, 2016; lizards: Muñoz-Mendoza et al. 2017). As such, we expect that the number of recognized species of *Virilastacus* will increase in the next few years. Finally, once the species richness of *Virilastacus* is adequately understood, a characterization of the genetic variation of the species of *Virilastacus* is also key to identifying intraspecific lineages that, in turn, should be also protected (Moritz, 1994). This knowledge is essential to maximize the maintenance of the evolutionary singularity, accurately evaluate conservation status, and design efficient conservation programs for Chilean freshwater crayfish, which are distributed in fragile habitats that are intensively being impacted by human intervention and climatic change.

## Acknowledgments

We thank Marcial Quiroga, Sara Rodriguez, Jaemy Romero, Richard Cadenillas, Andres Parada, and Martin Caicedo for their help during fieldwork and Alex Gonzalez for his help with laboratory work. Financial support was provided by the funding agencies SENESCYT and ANID (LA), FONDECYT Postdoctoral 3201059 (CME), and FONDECYT Regular 1221115 (GD).

## Statements and Declarations

The authors report there are no competing interests to declare.

